# Mycobacterial cell division arrest and smooth-to-rough envelope transition using CRISPRi-mediated genetic repression systems

**DOI:** 10.1101/2025.07.12.664508

**Authors:** Vanessa Point, Wafaa Achache, Janïs Laudouze, Eliana Sepulveda Ramos, Mickaël Maziero, Céline Crauste, Stéphane Canaan, Pierre Santucci

**Author notes:** Correspondence address to Pierre Santucci. Contributed equally as co-first authors (Vanessa Point, Wafaa Achache, Janïs Laudouze, Eliana Sepulveda Ramos).

## Abstract

The genetic basis underlying non-tuberculous mycobacteria (NTM) pathogenesis remains poorly understood. This gap in knowledge has been partially filled over the years through the generation of novel and efficient genetic tools, including the recently developed CRISPR interference (CRISPRi) technology. Our group recently capitalized on the well-established mycobacteria-optimized dCas9_Sth1_-mediated gene knockdown system to develop a new subset of fluorescence-based CRISPRi vectors that enable simultaneous controlled genetic repression and fluorescence imaging. In this Research Protocol, we use the model organism *Mycobacterium smegmatis* (*M. smegmatis*) as surrogate for NTM species and provide simple procedures to assess CRISPRi effectiveness. We describe how to evaluate the efficacy of gene-silencing when targeting essential genes but also genes involved in smooth-to-rough envelope transition, a critical feature in NTM pathogenesis. This protocol will have a broad utility for mycobacterial functional genomics and phenotypic assays in NTM species.

## Introduction

Infections caused by non-tuberculous mycobacteria (NTM) constitute an emerging global health concern, with the latest epidemiologic studies reporting an increase in cases worldwide [1–3]. NTM species are environmental organisms that primarily live in soil and aquatic niches [4]. Among them, only a few subsets are successful opportunist human pathogens that are responsible for a wide range of clinical syndromes from skin and soft-skin infections to potentially deadly pulmonary diseases [4].

NTM infections are notoriously difficult to diagnose and subsequently treat, especially species belonging to the *Mycobacterium avium* and *Mycobacterium abscessus* complex [5, 6]. Natural intrinsic resistance of NTM towards a large number of clinically available antibiotics, including anti-tuberculous drugs, make their treatment very challenging [6, 7]. Currently, a daily multi-drug regimen composed of 3-4 antibiotics is recommended, with treatment duration that can be very lengthy, often lasting 6 to 24 months, with poor clinical outcomes [5, 6]. The rising incidence of NTM infections in addition to the very limited number of efficient treatments available underscore the urgent need for new therapeutic approaches and improved public health strategies [8, 9].

To tackle this challenge, basic research represents a stepping stone by enabling a better understanding of the molecular mechanisms involved in NTM pathogenesis and the identification of new therapeutic targets. To do so, our research community has developed innovative functional genomics approaches coupled with sophisticated quantitative phenotypic assays. Over the years, the improvement of such technologies, especially the development of fast and efficient genetic tools in NTM [10–15], has allowed to identify and subsequently characterise critical genes and pathways that are either essential for bacterial growth [16–18] or required for the complete virulence in multiple hosts [19].

Accordingly, several genetic tools have been recently developed to selectively repress the expression of these specific drug targets in mycobacteria [16, 20–22]. Among them, we selected *rpoB* that encodes the β subunit of the RNA polymerase, as RpoB is targeted by compounds belonging to the rifamycin family, a critical class of drugs in the treatment of mycobacterial infections, including tuberculosis and certain NTM infections [23–25]. We also selected the *mmpL3* gene that encodes a critical transporter responsible for shuttling trehalose monomycolates to the mycobacterial cell wall and has been recently identified as a promising drug target [26–29].

In addition to identifying and targeting essential genes, the characterisation of critical steps required for optimal pathogenesis and virulence is important for developing tomorrow’s alternative therapeutic approaches. One of the hall-marks of NTM infection resides in their ability to irreversibly change the composition of their cell-wall, which during the infection process leads to an increased virulence [30–32]. This is particularly true for species from the *Mycobacterium avium* and *Mycobacterium abscessus* complex for which the transition from smooth to rough morphotypes has a significant implication for pathogenesis [4, 32, 33]. This modification, which is primarily linked to changes in the composition of the mycobacterial outer membrane, particularly the reduction or complete loss of glycopeptidolipids (GPL) [33–36], triggers the formation of large extracellular serpentine cords. These latter inhibit phagocytosis, prevent some of the host’s innate immune responses and exacerbate inflammation and cell death, therefore leading to the formation of abscesses or necrotic lung lesions [37, 38]. Understanding this transition is crucial for developing more appropriate therapies and, consequently improving the management of NTM infections. Therefore, we recently generated an anhydrotetracycline (ATc)-inducible mycobacterial hypomorph for which we can selectively repress the expression of *mmpL4b*, an essential component of the GPL translocase in NTM [22]. Such system allows to reversibly control the transition from smooth to rough, therefore constituting an interesting tool for the scientific community.

In this research protocol, we describe some simple methods to assess the efficiency of CRISPRi in *M. smegmatis* as a model organism to study NTM. We report procedures to evaluate gene-silencing effectiveness, focusing on both essential genes and those related to the smooth-to-rough transition of the bacterial envelope, an important aspect of NTM pathogenesis. By providing genetic tools, innovative experimental strategies and frameworks for better understanding the molecular and cellular functioning of these targets, we aim at potentially helping in the validation of new chemical entities.

## Materials

### Bacterial strains, storage and culture

- Recombinant *M. smegmatis* mc^2^155 strains harbouring plasmids pJL31 (Addgene #227428), pJL32 (Addgene #227429), and its derivatives pJL35, pJL36, pJL37 targeting *rpoB*, *mmpL3* and *mmpL4b* respectively [22].
- Complete Middlebrook 7H9 broth consisting of Middlebrook 7H9 base (BD Difco, Cat#271310) supplemented with 0.2% glycerol (Euromedex, Souffel-weyersheim, France, Cat#EU3550) and 0.05% Tween-80 (Sigma-Aldrich, Saint-Quentin Fallavier, France, Cat#P1754).
- Kanamycin sulfate (Euromedex, Cat#UK0010D) stock solution dissolved at 50 g.L^-1^ in distilled water.
- Glass Erlenmeyer 50 mL (Grosseron, Coueron, France, Cat#431.3332).
- Conventional pre-autoclaved 1.5 mL flat cap microcentrifuge plastic tubes (StarLab Tube One®, Orsay, France).
- Autoclaved 100% glycerol solution (Euromedex, Souffel-weyersheim, France, Cat#EU3550).

### Evaluating CRISPRi effectiveness on mycobacterial growth by microdilution or by serial dilution spot assays

- Recombinant *M. smegmatis* mc^2^155 strains harbouring plasmids pJL32 (Addgene #227429) and its derivatives pJL35 and pJL36 targeting *rpoB* and *mmpL3* respectively [22].
- Complete Middlebrook 7H9 broth consisting of Middlebrook 7H9 base (BD Difco, Cat#271310) supplemented with 0.2% glycerol (Euromedex, Souffel-weyersheim, France, Cat#EU3550) and 0.05% Tween-80 (Sigma-Aldrich, Saint-Quentin Fallavier, France, Cat#P1754).
- Complete Middlebrook 7H10 agar consisting of Middlebrook 7H10 agar base (BD Difco, Cat#262710) supplemented with 0.5% glycerol (Euromedex, Souffel-weyersheim, France Cat#EU3550).
- Square Petri Dishes 120 x 120 mm (Corning-Gosselin, Borre, France, Cat#BP124-05)
- Kanamycin sulfate (Euromedex, Cat#UK0010D) stock solution dissolved at 50 g.L^-1^ in distilled water.
- Hygromycin B (Toku-E, Sint-Denijs-Westrem, Belgium; #H007) stock solution dissolved at 50 g.L^-1^ in distilled water.
- Anhydrotetracycline (ATc) (Sigma-Aldrich, Saint-Quentin Fallavier, France; Supelco, Cat#37919) stock dissolved 1g.L^-1^ in ethanol (VWR, Rosny-sous-Bois, France, Cat#20823.362).
- Glass Erlenmeyer 50 mL (Grosseron, Coueron, France, Cat#431.3332)
- 50 mL sterile conical tubes in clarified polypropylene (Corning-Falcon, Avon, France, Cat#352070).
- 96-well flat-bottom Nunclon Delta Surface microplates with lid (Thermo-Fisher Scientific, Illkirch, France, Cat#167008).
- Spectrophotometer (Eppendorf Biophotometer, Montesson, France)
- Spectrophotometry cuvette ClearLine® (Dutscher, Bernolsheim, France, Cat#030101).
- TECAN Spark 10M Microplate reader (Tecan Group Ltd., Männedorf, Switzerland).
- ChemiDoc^TM^ MP Imaging System (Bio-Rad, Marnes-la-Coquette, France).
- Image Lab software version 6.1.0 (Bio-Rad, Marnes-la-Coquette, France).

### Monitoring CRISPRi-mediated smooth to rough morphotypes by macroscopic visualisation

- Recombinant *M. smegmatis* mc^2^155 strains harbouring plasmids pJL31 (Addgene #227428), pJL32 (Addgene #227429) and its derivative pJL37 targeting *mmpL4b* [22].
- Complete Middlebrook 7H9 broth consisting of Middlebrook 7H9 base (BD Difco, Cat#271310) supplemented with 0.2% glycerol (Euromedex, Souffel-weyersheim, France, Cat#EU3550) and 0.05% Tween-80 (Sigma-Aldrich, Saint-Quentin Fallavier, France, Cat#P1754).
- Complete Middlebrook 7H10 agar consisting of Middlebrook 7H10 agar base (BD Difco, Cat#262710) supplemented with 0.5% glycerol (Euromedex, Souffel-weyersheim, France Cat#EU3550).
- Round Petri Dishes 94 mm diameter (Greiner Bio-One, Les Ulis, France, Cat#633180).
- Kanamycin sulfate (Euromedex, Cat#UK0010D) stock solution dissolved at 50 g.L^-1^ in distilled water.
- Anhydrotetracycline (ATc) (Sigma-Aldrich, Saint-Quentin Fallavier, France; Supelco, Cat#37919) stock dissolved 1 g.L^-1^ in ethanol 96% (VWR, Rosny-sous-Bois, France Cat#20823.362).
- 50 mL Glass Erlenmeyer (Grosseron, Coueron, France, Cat#431.3332).
- 50 mL sterile conical tubes in clarified polypropylene (Corning-Falcon, Avon, France, Cat#352070).
- 10 µL sterile inoculation loops (Stirilab, Ružomberok, Slovakia, Cat#390530)
- ChemiDoc^TM^ MP Imaging System (Bio-Rad, Marnes-la-Coquette, France).
- Image Lab software version 6.1.0 (Bio-Rad, Marnes-la-Coquette, France).
- Fiji open-source software version 2.16.0 (https://imagej.net/software/fiji/) [39].
- Portable Digital USB 8 LED Mini-Microscope (Bysameyee) & a free compatible application such as OTG View2 Android App version 4.3.9.

### Quantifying glycopeptidolipid (GPL) levels during CRISPRi-mediated silencing of its translocase *mmpL4b*

#### Bacterial Culture, Sample Collection, Lyophilisation and Normalisation

- Recombinant *M. smegmatis* mc^2^155 strains harbouring plasmids pJL32 (Addgene #227429) and its derivative pJL37 targeting *mmpL4b* [22].
- Complete Middlebrook 7H9 broth consisting of Middlebrook 7H9 base (BD Difco, Cat#271310) supplemented with 0.2% glycerol (Euromedex, Souffel-weyersheim, France, Cat#EU3550) and 0.05% Tween-80 (Sigma-Aldrich, Saint-Quentin Fallavier, France, Cat#P1754).
- 50 mL Glass Erlenmeyer (Grosseron, Coueron, France, Cat#431.3332).
- 250 mL Glass Erlenmeyer (Grosseron, Coueron, France, Cat#431.3472).
- Kanamycin sulfate (Euromedex, Cat#UK0010D) stock solution dissolved at 50 g.L^-1^ in distilled water.
- Anhydrotetracycline (ATc) (Sigma-Aldrich, Saint-Quentin Fallavier, France, Supelco, Cat#37919) stock dissolved 1 g.L^-1^ in ethanol 96% (VWR, Rosny-sous-Bois, France, Cat#20823.362).
- 50 mL sterile conical tubes in clarified polypropylene (Corning-Falcon, Avon, France, Cat#352070).
- Phosphate Buffer Saline (10 mM phosphate buffer, and 3 mM KCl, pH 7.4) (Merck-Millipore, Darmstadt, Germany, Cat#524650)
- Spectrophotometer (Eppendorf Biophotometer, Montesson, France)
- Centrifuge 5810R (Eppendorf, Montesson, France)
- Lyophilizator Alpha 1-4 LSCbasic (Christ, Osterode am Harz, Germany)

#### Lipids Extraction & Sample preparation

- Centrifuge 5810R (Eppendorf, Montesson, France).
- Rotary evaporator Hei-VAP Advantage (Heidolf Instruments, Schwabach, Germany).
- Nitrogen flow with Nitrogen 4.5 (Linde, Lyon, France, Cat#UN1066).
- CHCl_3_ (VWR, Rosny-sous-Bois, France, Cat#22711.290).
- CH_3_OH (VWR, Rosny-sous-Bois, France, Cat#20847.318).
- Glass Pasteur pipette 150 mm (Dutscher, Bernolsheim, France, Cat#065421)
- 100 mL Glass bottle with ground-glass neck (Grosseron, Coueron, France, Cat#4361402).
- 15 mL Glass funnel (Dutscher, Bernolsheim, France, Cat#068958).
- 2 mL amber glass vial (Merck, Darmstadt, Germany, Cat#27000).
- Ashless filter paper Grade 40 (Whatman®, Cytiva, Saint-Germain-en-Laye, France, Cat# 1440-090).
- 100 mL piriform round-bottom flask (Grosseron, Coueron, France, Cat#9.012024).
- 50 mL sterile conical tubes in clarified polypropylene (Corning-Falcon, Avon, France, Cat#352070).

#### Thin Layer Chromatography (TLC) running and development

- Semi-automatic sample dispenser Linomat 5 (CAMAG, Muttenz, Switzerland, Cat#022.7808).
- 100 µL syringe for Linomat 5 (CAMAG, Muttenz, Switzerland, Cat#695.0014).
- Nitrogen bottle with pressure gauge set at 4-5 bar (Linde, Lyon, France, Cat#UN1066).
- TLC Silica gel 60 F254 100*200 mm (Merck, Darmstadt, Germany, Cat#1.05729.0001).
- Automatic Developing Chamber 2 – ADC2 (CAMAG, Muttenz, Switzerland, Cat#022.8380) coupled with humidity control (CAMAG, Muttenz, Switzerland, Cat#022.8360) using MgCl_2_ (Sigma-Aldrich, Saint-Quentin Fallavier, France, Cat#13152) 33% salt solution in distilled water.
- *visionCATS* software basic version (CAMAG, Muttenz, Switzerland, Cat#028.0000).
- Filter paper for chamber saturation (CAMAG, Muttenz, Switzerland, Cat#022.8370).
- Flat bottom chamber with glass lid for 200*200 mm plates (CAMAG, Muttenz, Switzerland, Cat#022.5250).
- TLC plate heater III (CAMAG, Muttenz, Switzerland, Cat#022.3306).
- TLC visualizer II (CAMAG, Muttenz, Switzerland, Cat#022.9811).
- Glass reagent sprayer 100 mL (CAMAG, Muttenz, Switzerland, Cat#022.6100).
- 10 µL glass capillaries (CAMAG, Muttenz, Switzerland, Cat#022.7730).
- TLC spray cabinet with fan (CAMAG, Muttenz, Switzerland, Cat#022.6230).
- Anthrone (Sigma-Aldrich, Saint-Quentin Fallavier, France, Cat#319899).
- H_2_SO_4_ 95% (VWR, Rosny-sous-Bois, France, Cat#20685.295).
- CH_3_CH_2_OH 96% (VWR, Rosny-sous-Bois, France, Cat#20823.362).
- CHCl_3_ (VWR, Rosny-sous-Bois, France, Cat#22711.290).
- CH_3_OH (VWR, Rosny-sous-Bois, France, Cat#20847.318).
- MgCl_2_.6H_2_O (Sigma-Aldrich, Saint-Quentin Fallavier, France, Cat#13152).
- 100 mL Glass flask (Grosseron, Coueron, France, Cat#4361402).

#### Quantitative Analysis of GPL Levels

- ChemiDoc^TM^ MP Imaging System (Bio-Rad, Marnes-la-Coquette, France).
- Image Lab software version 6.1.0 (Bio-Rad, Marnes-la-Coquette, France).
- Fiji open-source software version 2.16.0 (https://imagej.net/software/fiji/) [39].
- Microsoft office LTSC Standard 2021 including Excel (version 2018).
- R Studio software (The R Project for Statistical Computing), with the ggplot2 package (version 3.5.1).

## Methods

### Generation, Storage and Maintenance of Mycobacterial CRISPRi Strains

*Note: Vectors from the pIRL and pJL series used to perform CRISPRi are available on Addgene* (https://www.addgene.org/)*. The generation of the plasmids described in this research protocol has been described previously* [22] *following the well-established CRISPRi cloning protocol reported by Wong et al.* [40]*. Once generated and sequenced, CRISPRi vectors have been electroporated in M. smegmatis mc^2^ 155 following the procedure described by Goude et al.* [41]*. Recombinants clones have been selected, further expanded, and frozen glycerol stocks of each genetic background have been stored at -80°C and are subsequently used in this study*.

#### Initiating mycobacterial starter cultures from frozen stocks

1. Thaw pre-aliquoted 500 µL-1 mL glycerol stocks of recombinant *M. smegmatis* mc^2^155 strains harbouring plasmids pJL31, pJL32, pJL35, pJL36 or pJL37 according to your experimental plan.
2. Prepare 20 mL of complete Middlebrook 7H9 broth supplemented with kanamycin at a final concentration of 50 mg.L⁻¹ for each bacterial background in pre-autoclaved 50 mL Erlenmeyer flasks.
3. Inoculate the entire content of each glycerol stock tubes within the 20 mL of complete medium.
4. Incubate the cultures at 37°C under shaking at 180 rpm for 48-72 hours.
5. After 48-72 hours, collect a 1 mL aliquot of each saturated bacterial culture.
6. Dilute 100 µL of each bacterial culture into 1000 µL final of complete Middlebrook 7H9 broth and transfer this into a spectrophotometry cuvette.
7. Evaluate bacterial growth spectrophotometrically by measuring optical density (OD_600nm_) using complete Middlebrook 7H9 broth only as blank.

#### Subculturing of mycobacterial cultures

8. Prepare 50 mL of complete Middlebrook 7H9 broth supplemented with kanamycin at a final concentration of 50 mg.L⁻¹ for each bacterial background in pre-autoclaved 250 mL Erlenmeyer flasks.
9. Subculture each strain according to the recorded OD_600nm_ of the starter cultures by diluting them to obtain an initial OD_600nm_ comprised between 0.025-0.05.
10. Incubate the cultures at 37°C under shaking at 180 rpm for 16-24 hours.
11. After 16-24 hours, evaluate bacterial growth spectrophotometrically by measuring optical density (OD_600nm_) as described above.

*Note: M. smegmatis mc^2^ 155 has an approximate doubling time of 4 hours in rich medium; therefore, 16-24 hours of incubation usually enable to have exponentially growing bacterial cultures with OD_600nm_ comprised between 0.5 and 1.5*.

*Note: At that stage, exponentially growing cultures can be used to perform a wide range of phenotypic assays, such as evaluating CRISPRi-mediated inhibition of mycobacterial growth by microdilution or by serial dilution spot assays, but also performing smooth-to-rough transition visual assays*.

### Evaluating CRISPRi effectiveness on mycobacterial growth by microdilution or by serial dilution spot assays

#### 1. Monitoring essential genes targeting by CRISPRi using microdilution assays

##### A. Microplate inoculation using standardized bacterial inoculum

1. Using exponentially growing cultures as described below, prepare bacterial inoculum for each recombinant strain to be tested by adjusting the OD_600nm_ to 0.005. In this experiment, we will use pJL32, pJL35 and pJL36 *M. smegmatis* mc^2^ 155 recombinant strains. Assuming that an OD_600nm_ of 1 corresponds to approximately to 1 × 10^8^ CFU.mL^-1^, an inoculum of OD_600nm_ 0.005 will result in a bacterial concentration of approximately 5 × 10^5^ CFU.mL^-1^.
2. Within a 96-well plate, perform two-fold serial dilutions of ATc ranging from 200 ng.L⁻¹ to 0.08 ng.L⁻¹ in 100 µL of media (Fig 1A).
3. Add 100 µL of the bacterial inoculum to each well, ensuring a final volume of 200 µL per well and final ATc concentration ranging from 100 ng·L⁻¹ to 0.04 ng·L⁻¹ (Fig 1A).
4. Include all the required control conditions, such as a positive growth control (bacterial inoculum in complete media without ATc and/or with vehicle only), a negative sterility control (complete Middlebrook 7H9 broth without inoculum) and a positive control of bacterial growth inhibition (bacterial inoculum with complete Middlebrook 7H9 broth containing 50 mg·L⁻¹ of Hygromycin B) (Fig 1A).
5. Include a technical triplicate in the plate for each condition (Fig 1A).
6. Measure the OD_600nm_ using the 96-well microplate reader to assess the OD_600nm_ at timepoint 0h.

*Note: At this stage the OD_600nm_ is similar between inoculated media and sterility control condition, therefore leading a starting OD_600nm_ corrected value of 0*.

7. Incubate the microplates at 37°C without shaking, preferably in a humidity chamber to prevent evaporation.

**Figure 1.**
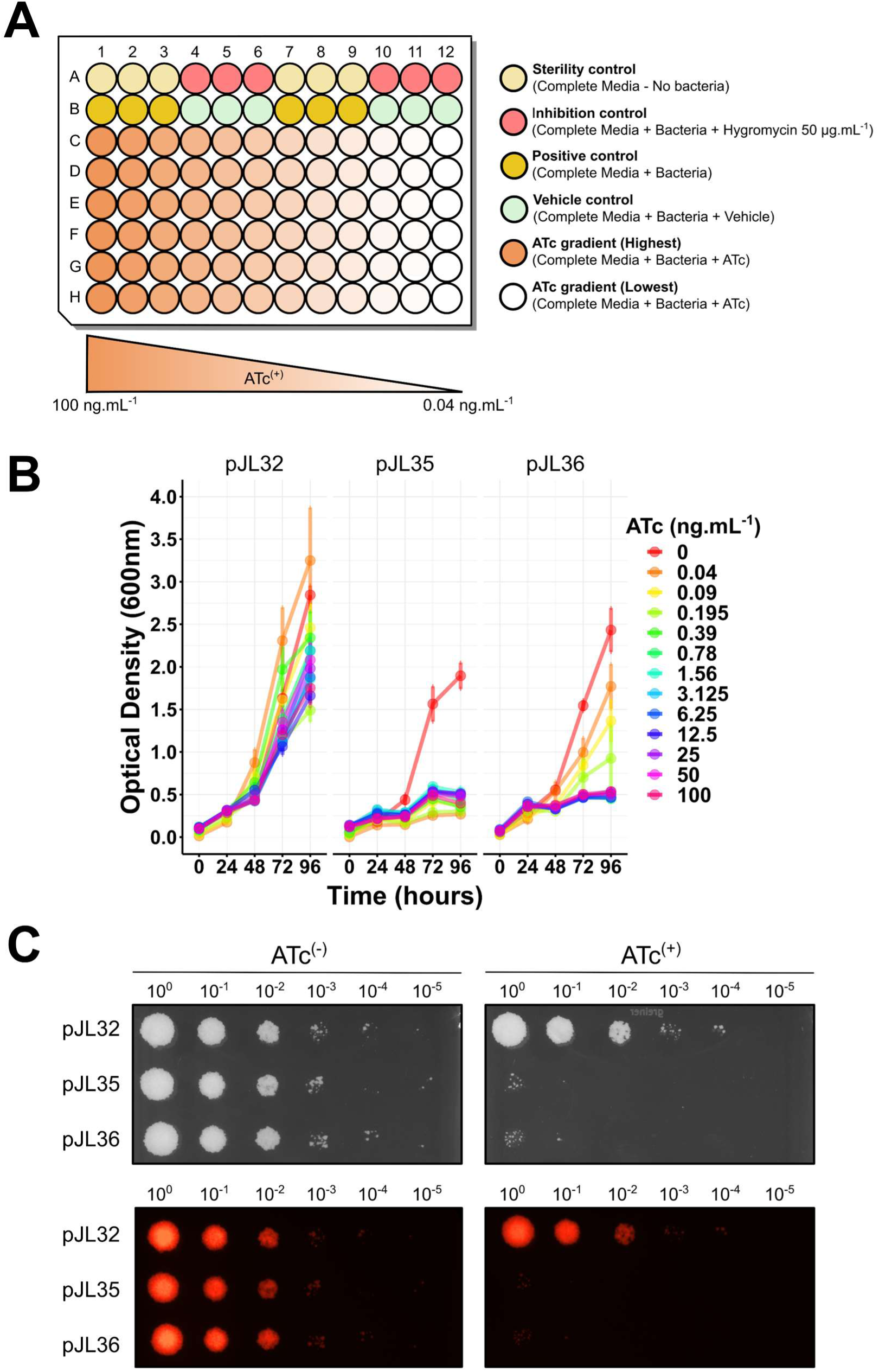
Monitoring of CRISPRi effectiveness on mycobacterial growth while targeting essential genes. **(A)** Schematic representation of a standard microplate used to perform a microdilution-based assessment of CRISPRi effectiveness on mycobacterial growth. This representation is suitable for testing two strains within the same microplate. **(B)** Effect of CRISPRI-mediated gene silencing on *M. smegmatis* mc^2^ 155 growth in liquid media. Growth profiles of pJL32, pJL35 and pJL36 recombinant strains were assessed by measuring optical density at 600nm, and performed in the absence or in the presence of increasing concentrations of ATc ranging from 0 to 100 ng.mL^-1^. An individual colour code has been used to identify each concentration. Of note, while the inhibition is almost complete, all the conditions are displayed as superimposed on the graph. **(C)** Effect of CRISPRI-mediated gene silencing on *M. smegmatis* mc^2^ 155 growth on agar media. Serial dilutions of *M. smegmatis* mc^2^ 155 recombinant strains pJL32, pJL35 and pJL36 were spotted onto 7H10 agar media either in the absence (left panel) or presence (right panel) of 100 ng.mL^-1^ of ATc. Bright light (top) and their corresponding red fluorescent profiles (bottom) are shown.

##### B. Recording of CRISPRi-mediated bacterial growth inhibition and data analysis

8. Monitor bacterial growth by measuring the OD_600nm_ every 24 hours using a 96-well plate spectrophotometer.
9. Repeat this operation for 4 consecutive days to obtain data at different timepoints corresponding to 0 h, 24 h, 48 h, 72 h and 96 h.
10. At each stage, ensure that all the control conditions match the expected results.
11. Subsequently collect and organise all the data using Microsoft Excel or any alternative software. This step consists of preparing a matrix summarizing the data, including all replicates, the recorded OD_600nm_, and the corrected OD_600nm_ values obtained by subtracting the blank.
12. Export the results summarized in this matrix as “.csv” file and perform data analysis using R studio or any other data visualisation software that can display line plots (Fig 1B).
13. Display bacterial growth dynamics for each strain in the absence or the presence of increasing concentrations of ATc by generating the corresponding plots for each strain. Representative results obtained with pJL32, pJL35 and pJL36 are displayed in Fig 1B.

*Note: This whole workflow is repeated two or three times in order to get more biological replicates and make sure that the observed phenotypes are reproductible over independent experiments*.

#### 2. Monitoring essential genes targeting by CRISPRi using agar spot assays

##### A. Preparation of agar plates

*Note: Prepare 7H10 agar media and autoclave it according to the manufacturer instructions. We usually prepare 5 L batches that are aliquoted in twelve independents 500 mL glass bottles containing approximately 400 mL of media*.

1. Use a conventional microwave to melt a 500 mL glass bottle of complete 7H10 media.
2. Once the liquid is homogeneous, leave the bottle cool down in 50-55°C water bath for at least 20-30 min.
3. Once at the appropriate temperature, add kanamycin stock solution to the media to a final concentration of 50 mg. L^-1^ and vigorously shake to homogenise.
4. Use half of this bottle of media to dispense 50 mL of medium into square petri dishes. This batch of plates will constitute the uninduced control condition also referred to as ATc^(-)^.
5. Use the remaining half to add ATc at final concertation of 100 ng.mL^-1^.
6. Dispense 50 mL of medium into 120x120 mm square petri dishes. This batch of plates will constitute the induced condition also referred to as ATc^(+)^.
7. Leave the plates open for a few minutes under a Bunsen burner or microbiological safety cabinet in order to let them solidify and dry.
8. Once dry, the plates can be used directly or store at 4°C for later use.

##### B. Preparation of bacterial inoculum and dropping onto agar plates

9. Using exponentially growing cultures, prepare bacterial inoculum for each recombinant strain by adjusting the OD_600nm_ to 0.1. In this experiment, we will use pJL32, pJL35 and pJL36 *M. smegmatis* mc^2^ 155 recombinant strains. Assuming that an OD_600nm_ of 1 corresponds to approximately to 1 × 10^8^ CFU.mL^-1^, this inoculum will result in a bacterial concentration of approximately 1 × 10^7^ CFU.mL^-^_1._
10. Using this inoculum at OD_600nm_ 0.1, perform 10-fold serial dilution by collecting 20 μL of the inoculum and mix it with 180 μL of complete Middlebrook 7H9 broth medium. This can be done in conventional 1.5 mL sterile microcentrifuge plastic tubes or in 96-well plates.
11. Repeat this operation 5 to 6 times by changing tips for each dilution. Following this procedure, you should reach a desired dilution ranging from 10^0^ to 10^-5^ or 10^-6^ corresponding to a theoretical number of 1 × 10^2^ CFU.mL^-1^ or 1 × 10^1^ CFU.mL^-1^ respectively.
12. Using a single or multi-channel pipette, collect 10 μL of each dilution and carefully place it on a 7H10 agar plate containing 50 mg.L^-1^ of kanamycin in the absence or presence of 100 ng.mL^-1^ of ATc.
13. Once the drops have been dispensed, let the plate dry for 10-20 minutes. Do not move the plate as this can result in altered morphology and/or mixing of the drops.
14. When dry, incubate the plates at 37°C for 3-4 days.
15. Scanning of each plate is performed using ChemiDoc^TM^ MP Imaging System coupled with the Image Lab software. Bright light acquisition is performed by using the ‘White Epi Illumination’ setting combined with the ‘Standard Filter’, while red fluorescence acquisition is performed by using the ‘Green Epi Illumination’ setting combined with the ‘605/50 nm Filter’.
16. If required, individual raw images can be exported from Image Lab as TIFF files 600 dpi and displayed with the open-source software Fiji for further processing. Results obtained with the pJL32, pJL35 and pJL36 strains are displayed in Fig 1C and are representative of 2 independent biological replicates.

### Monitoring CRISPRi-mediated smooth-to-rough morphotype transition by macroscopic visualisation

*Note: This step describes two-distinct approaches to evaluate CRISPRi-mediated changes in morphotypes. Both approaches have been previously reported here* [22].

1. Using exponentially growing cultures of pJL31, pJL32 and pJL37, prepare bacterial inoculum for each recombinant strain by adjusting the OD_600nm_ to 0.1. Assuming that an OD_600nm_ of 1 corresponds to approximately to 1 × 10^8^ CFU.mL^-1^, this inoculum will result in a bacterial concentration of approximately 1 × 10^7^ CFU.mL^-^_1._
2. Using pJL32 and pJL37 inoculum at OD_600nm_ 0.1, drop 10 µL on 7H10 agar media containing 50 mg.L^-1^ of kanamycin in the absence or presence of 100 µg. L^-1^ of ATc as described previously.
3. Subsequently, streak the drop using an inoculation loop following the established quadrant technique.
4. Incubate the plates at 37°C for 2-4 days.
5. Scan each plates using ChemiDoc^TM^ MP Imaging System coupled with the Image Lab software. Bright light acquisition is performed by using the ‘White Epi Illumination’ setting combined with the ‘Standard Filter’, green fluorescence acquisition is performed by using the ‘Blue Epi Illumination’ setting combined with the ‘530/28 nm Filter’ and red fluorescence acquisition is performed by using the ‘Green Epi Illumination’ setting combined with the ‘605/50 nm Filter’.
6. If required, individual raw images can be exported from Image Lab as TIFF files 600 dpi and displayed with the open-source software Fiji for further processing. Results are displayed in Fig 2A.
7. Alternatively, plating can be performed on the same Petri dish by mixing multiple inoculums expressing different fluorophores as previously described [22]. A representation of a mixed experiment with pJL31 and pJL37 is reported in Fig 2B-E.
8. Finally, higher magnification pictures of colonies of interest can also be visualized using a portative digital microscope coupled with a basic smartphone. Results are displayed in Fig 2F.

**Figure 2.**
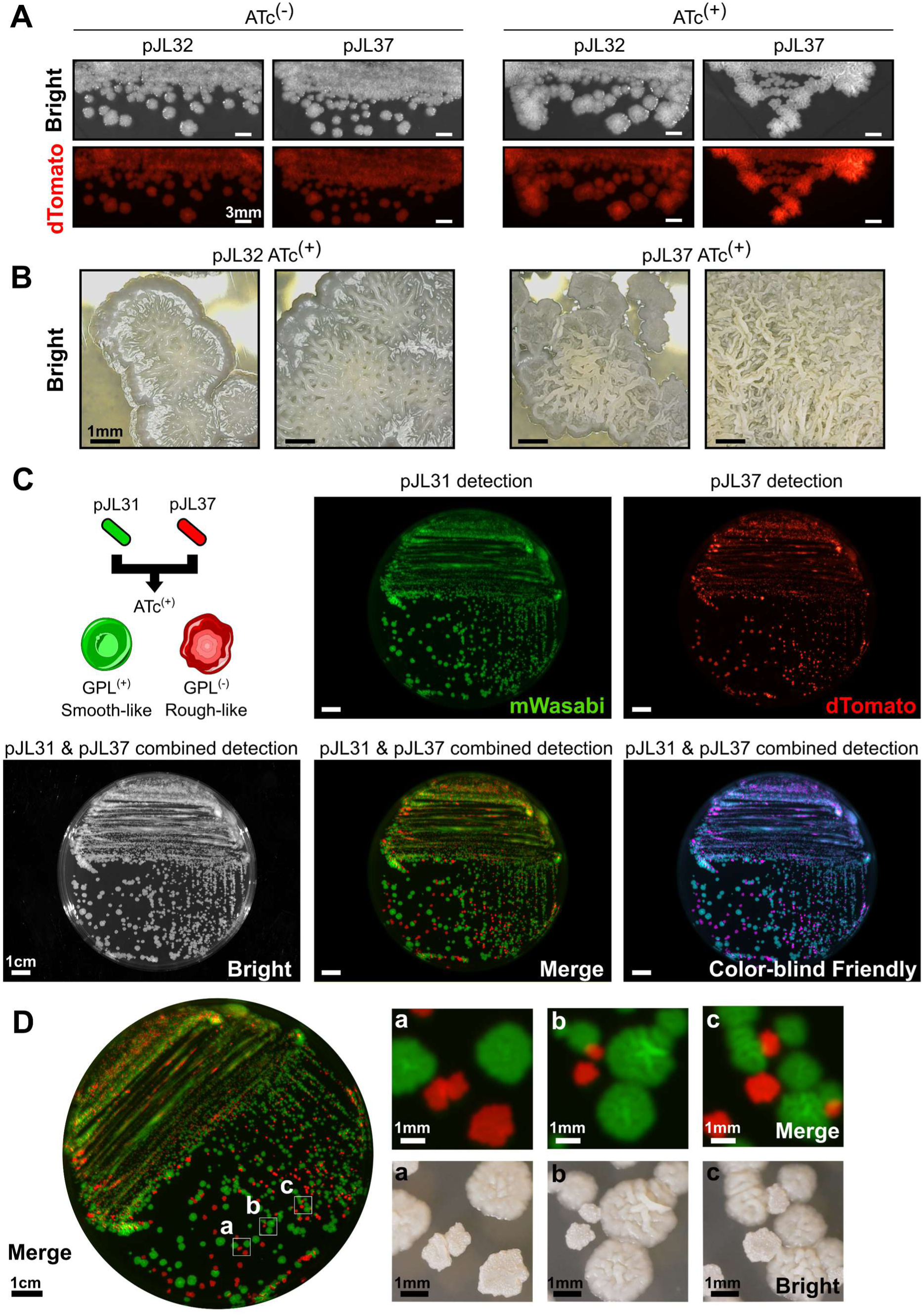
Phenotypic visualisation of smooth-to-rough transition using inducible CRISPRi-mediated silencing of GPL translocation. **(A)** Colony morphotype analysis of *mmpL4b* CRISPRi-mediated silencing using the pJL32 and pJL37 vectors. Cells were grown in 7H10 solid medium in the presence or absence of 100 ng.mL^−1^ of ATc for 72 h before being imaged using Chemidoc MP imaging system. Bright light (top) and their corresponding red fluorescent profiles (bottom) are shown. Scale bar represents 3 mm. **(B)** Close up analysis of smooth-like (pJL32) and rough-like (pJL37) morphotypes obtained on 7H10 solid medium in the presence 100 ng.mL^−1^ of ATc for 72 h with a portable digital camera. Smooth-like morphotype displays more regular and rounded edges with a shiny and glossy appearance while rough-like morphotype is characterized by more irregular edges and drier surface. Scale bar represents 1 mm. **(C)** Analysis of *M. smegmatis* mc^2^ 155 pJL31 and pJL37 morphologies and fluorescence profiles when mixed on solid medium. Approximately, 1.10^6^ CFU of *M. smegmatis* mc^2^ 155 pJL31 and 1.10^6^ CFU of *M. smegmatis* mc^2^ 155 pJL37 were mixed before being plated on 7H10 solid medium containing 100 ng.mL^−1^ of ATc. After 3–4 days at 37 °C, the plates were imaged using Chemidoc MP imaging system. Bright light, green fluorescence, red fluorescence, merge and a colourblind friendly merge micrographs are displayed. Scale bar represents 1 cm. **(D)** Zoom caption of the merge channel is displayed on the left, and selected area from (a), (b) and (c) have been further imaged using a portable digital microscope. Their corresponding micrographs are showed at the right of the panel. Scale bars represent 1 cm and 1 mm respectively.

*Note: At this stage, an additional reversion assay can be performed to confirm that the observed phenotypes are all due to the CRISPRi-mediated inducible system. Hence, some individual pJL31 and pJL37 smooth and rough colonies from* Fig2 *are randomly picked and used to inoculate 5 mL of ATc-free complete 7H9 Middlebrook media containing 50 mg.L^-1^ of kanamycin. Independent cultures are further incubated for 48-72 h, and the procedure described below is repeated on both ATc^(-)^ and ATc^(+)^ agar plates. Representative results obtained following this procedure are displayed in* Fig 3.

**Figure 3.**
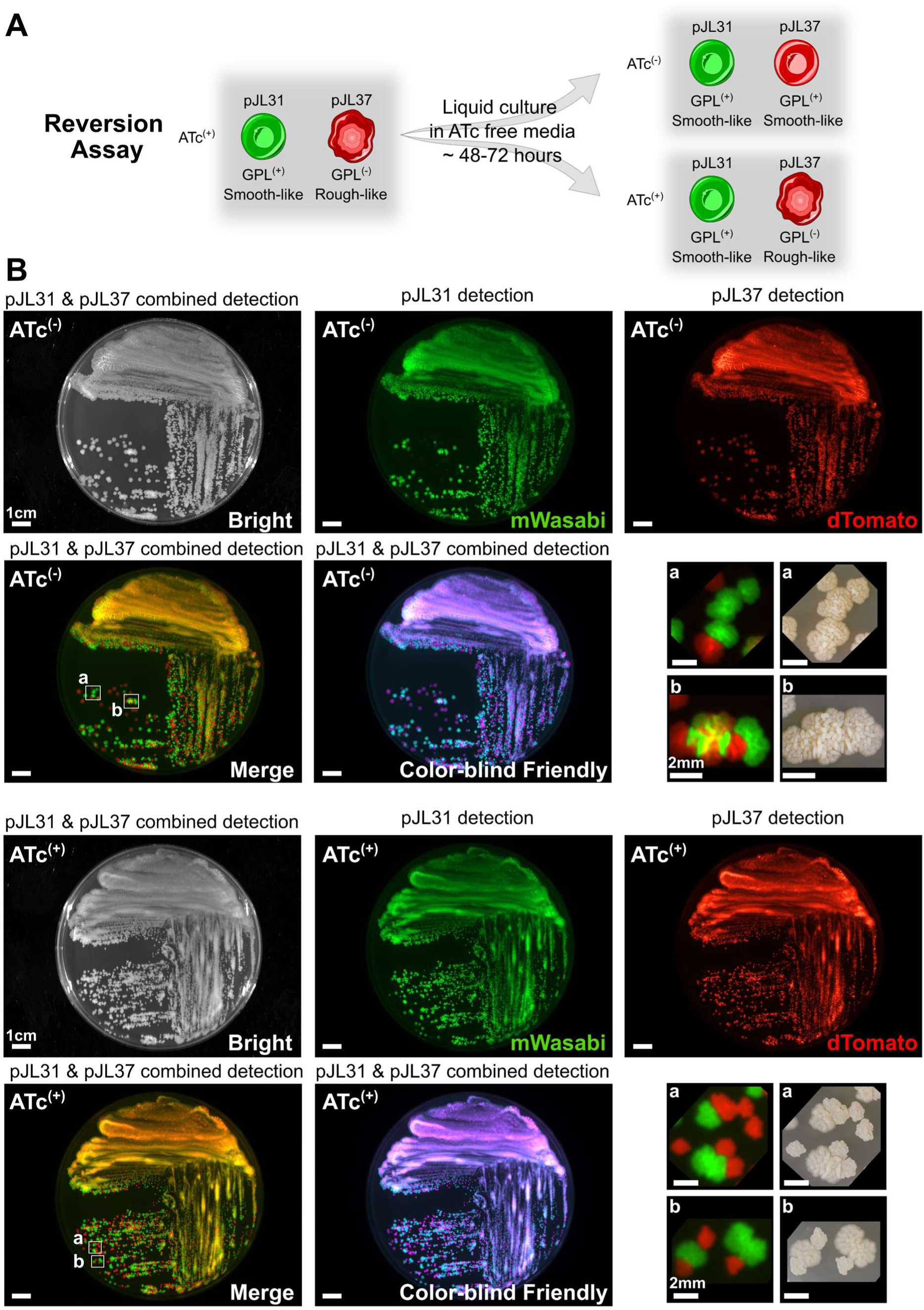
Assessing the reversible nature of the smooth-to-rough transition using inducible CRISPRi-mediated silencing of GPL translocation. **(A)** Representative scheme of the principle of the reversion assay performed in this study. **(B)** Analysis of *M. smegmatis* mc^2^ 155 pJL31 and pJL37 morphologies and fluorescence profiles when mixed on solid medium. Approximately, 1.10^6^ CFU of *M. smegmatis* mc^2^ 155 pJL31 and 1.10^6^ CFU of *M. smegmatis* mc^2^ 155 pJL37 collected from ATc-free liquid medium were mixed before being plated on 7H10 solid medium in the absence (top panels) or in the presence (bottom panels) of 100 ng.mL^−1^ of ATc. After 3–4 days at 37 °C, the plates were imaged using Chemidoc MP imaging system. Bright light, green fluorescence, red fluorescence, merge and a colourblind friendly merge micrographs are displayed. Scale bar represents 1 cm. Zoom captions of the merge channel, appear as selected areas (a) and (b) are displayed on the right. Corresponding colonies have been further imaged using a portable digital microscope, flipped and manually cropped to match the fluorescence images. Scale bars represent 1 cm and 2 mm respectively.

### Quantifying glycopeptidolipid (GPL) levels during CRISPRi-mediated silencing of its translocase *mmpL4b*

#### A. Subculturing of mycobacterial cultures and sample lyophilisation

1. Prepare 100 mL of complete Middlebrook 7H9 broth supplemented with kanamycin at a final concentration of 50 mg.L⁻¹ in pre-autoclaved 250 mL or 500 mL Erlenmeyer flasks. For the experiment presented in this manuscript, 9 Erlenmeyer flasks were required.
2. Subculture pJL32 and pJL37 strains according to recorded OD_600nm_ of the starter cultures by diluting them in order to obtain at an initial OD_600nm_ of 0.025-0.05.
3. Incubate pJL32 cultures at 37°C under shaking at 180 rpm for 48 hours in the absence or in the presence of 100 ng.mL^-1^ of ATc.
4. Incubate pJL37 cultures at 37°C under shaking at 180 rpm for 48 hours in the absence or in the presence of increasing concentrations of ATc ranging from 0 to 100 ng.mL^-1^ (*e.g.* 0, 0.1, 1, 5, 10, 20, 50 and 100 ng.mL^-1^).
5. After 48 hours, centrifuge 100 mL of bacterial cultures at 5000 rpm for 10 min in 50 mL conical tubes.
6. Wash the pellet twice using sterile PBS buffer pH7.4.
7. Lyophilize the bacterial pellets overnight using the Lyophilizator Alpha 1-4 LSCbasic.
8. Collect the lyophilized samples.
9. Weigh the same amount of dry material for all samples (between 100 and 150 mg) in 50 mL conical tube.

#### B. Conventional workflow for total mycobacterial lipid extraction

*Note: The following steps involve working with volatile organic solvents, and must therefore, be carried out in a fume cupboard or chemical hood*.

10. Add 30 mL of CHCl_3_/CH_3_OH (1:2, *v/v*) in each dry pellet and incubate on a rotating mixer for 24 hours at room temperature (RT).
11. Centrifuge at 5000 rpm for 10 min at RT.
12. Transfer the organic solvent (OS_1_) in a glass bottle using a glass Pasteur pipette.
13. Add 30 mL of CHCl_3_/CH_3_OH (1:1, *v/v*) to each residual pellet and incubate on a rotating mixer for 24 hours at room temperature (RT).
14. Centrifuge at 5000 rpm for 10 min at RT.
15. Transfer the organic solvent (OS_2_) and pool it with OS_1_ in a glass bottle.
16. Add 30 mL of CHCl_3_/CH_3_OH (2:1, *v/v*) to each residual pellet and incubate on a rotating mixer for 24 hours at room temperature (RT).
17. Centrifuge at 5000 rpm for 10 min at RT.
18. Transfer the final organic solvent (OS_3_) using a Pasteur pipette and pool it with OS_1_ and OS_2_ in the glass bottle.
19. Filter the mixed organic solvents (OS_1_, OS_2_ and OS_3_) into a 100 mL round-bottom flask through ash filter paper using a glass funnel.
20. Use a rotary evaporator to dry the mixed solvents until complete evaporation.
21. Resuspend the total extracted lipids in a minimum of CHCl_3_/CH_3_OH (2:1, *v/v*, ∼500 µL) and transfer the extract using a glass Pasteur pipette in a pre-tared vial.
22. Rinse twice the round-bottom flask with CHCl_3_/CH_3_OH (2:1, *v/v*, ∼500 µL) in order to make sure to recover all the lipids.
23. Evaporate to dry the lipid extracts in a vial under nitrogen flow.

*Note: It is recommended to weigh the dried lipids at this stage to estimate the quantity*.

24. After evaporation under nitrogen flow, resuspend the lipids in 800 µL of CHCl_3_/CH_3_OH (2:1, *v/v*) and shake vigorously for 1 min.

*Note: The resuspension volume depends on the weight of total lipids obtained. Generally, resuspend at a concentration of around 25 mg.mL^-1^*.

*Note: At that stage, GPL can be subsequently analyzed by TLC using either a conventional approach with manual loading or a dedicated CAMAG semi-automated equipment for improved sensitivity, resolution and reproducibility. Both methods are described below*.

#### C. TLC analysis without CAMAG equipment

25. Successively place 10 µL of samples on the TLC plate using a glass capillary or a pipette with 10 µL tip, ensuring to rinse the capillary thoroughly with CHCl_3_/CH_3_OH (2:1, *v/v*) or changing tip between each sample.
26. Introduce 100 mL of CHCl_3_/CH_3_OH (9:1, *v/v*) elution solvent into the Flat Bottom Chamber.
27. Place the TLC plate in the chamber and let the eluent migrate.
28. During the migration, prepare the anthrone solution, weigh 0.2 g of anthrone into a 100 mL glass flask and add 100 mL of ethanol containing 20% H_2_SO_4._
29. Remove the TLC from the chamber when the eluent has risen by approximately 8 cm.
30. Dry the TLC and position it in the TLC spray hood coupled with a fan.
31. Spray the entire plate evenly with anthrone solution.
32. Heat the plate for 3-5 min at 120°C. GPL species should appear blue on the TLC.
33. Perform imaging of the TLC (Fig 4A).

#### D. TLC analysis using semi-automated CAMAG equipment

34. Open the nitrogen bottle connected to the Linomat 5 sample applicator.
35. Switch on all the required equipment including the Linomat 5, ADC2 and Visualizer 2, then open the *visionCATS* software.
36. After preparing the analysis method and settings on *visionCATS*, place a TLC plate on the Linomat and launch the program.
37. Using the CAMAG syringe, successively dispense 7 µL of sample onto the TLC plate, by ensuring to rinse the syringe thoroughly with CHCl_3_/CH_3_OH (2:1, *v/v*) between each sample.

*Note: The visionCATS software allows to adjust various parameters with the Linomat 5 sample applicator such as deposit width or spacing between deposits for example, depending on the number of samples on the plate*.

38. Prepare 60 mL CHCl_3_/CH_3_OH (9:1, *v/v*) as elution solvent.
39. Place a filter paper in the ADC2 migration chamber and wet it with the elution solvent.
40. Introduce 10 mL of elution solvent into the “Development” module and 25 mL into the “Saturation” module.
41. Place the TLC in the ADC2 and run the program.

**Figure 4.**
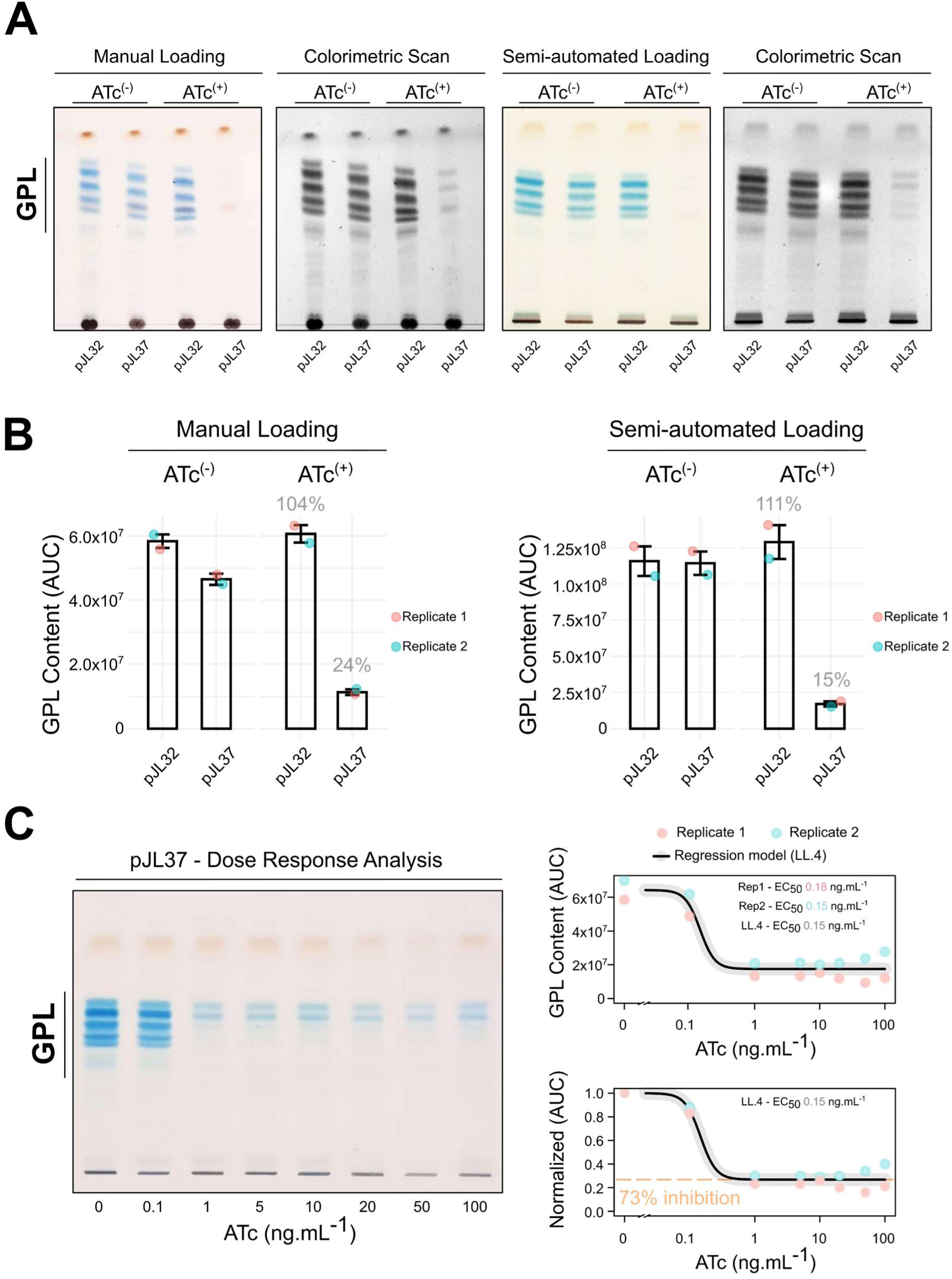
Quantitative analysis of GPL levels upon CRISPRi-based repression of *mmpL4b*-mediated export. **(A)** Representative TLC of GPL analysis of *M. smegmatis* mc^2^ 155 pJL32 or pJL37 in the presence or in the absence of ATc. Cells were grown in 7H9 broth ± 100 ng.mL^−1^ of ATc for 48 h. Lipid extraction from normalized dry pellets was analysed by TLC either by performing a manual or semi-automated loading. GPL appear in blue upon development and TLC plates are further scanned in black and white to perform densitometric analyses. **(B)** Corresponding quantitative analysis of GPL content of *M. smegmatis* mc^2^ 155 pJL32 or pJL37 grown in the presence or in the absence of ATc. Each TLC were performed in two technical replicates, shown in cyan and pink. GPL content was determined by quantifying the total area under the curve (AUC). Values indicated in grey illustrate the % of GPL remaining in comparison to the ATc^(-)^ condition. **(C)** Representative TLC and quantitative GPL analysis of *M. smegmatis* mc^2^ 155 pJL37 in the presence of increasing concentration of ATc. Cells were grown in 7H9 broth ± 0, 0.1, 1, 5, 10, 20, 50 or 100 ng.mL^−1^ of ATc for 48 h. Lipid extraction from normalized dry pellets was analysed by TLC as done in panel A and B using the semi-automated method. TLC were performed in two technical replicates, shown in cyan and pink. GPL content was determined by quantifying the total area under the curve (AUC). A four-parameter log-logistic regression (LL4) was applied to determine the effective concentration 50 (EC_50_) leading to 50% reduction in GPL content in between the two plateaus of the built model. The analysis was also performed by normalizing the AUC of the no ATc condition to 1 (Bottom panel). A maximal effect was observed reaching 73% of GPL reduction and is highlighted in orange.

*Note: The following settings were used to perform our experiments with the ADC2. Front solvent: 75 mm; Activation: 20 min; Saturation: 20 min; Drying: 5 min*.

42. During the migration, prepare the anthrone solution. Weigh 0.2 g anthrone into 100 mL glass flask and add 100 mL ethanol containing 20% H_2_SO_4_.
43. At the end of migration, remove the TLC plate from the ADC2 and position it in the TLC spray hood coupled with a fan. Spray the entire plate evenly with anthrone solution.
44. Heat the plate for 3-5 min at 120°C. GPL species should appear blue on the TLC.
45. Place the plate in the Visualizer 2 and take the respective picture under white light (Fig 4A).

*Note: GPL levels can be determined by semi-quantitative analysis using different equipment and software. The next section presents two methods: the first using ChemiDoc MP imager/Image Lab software and the second using the open-source Fiji software*.

#### E. Quantitative analysis using the ChemiDoc MP imager and Image Lab software

46. Switch on the ChemiDoc and open the Image Lab software.
47. Position the White Light Conversion Screen and place the TLC in the center.
48. Set and adjust the camera zoom.
49. Acquire the TLC image with the Colorimetric program using the “Blots” building block. Using this program, the GPL appear in black on a white background (Fig 4A).
50. In the Analysis Tool Box, select the “Lane and Bands” building block, and manually set the number of samples to be quantified in “Lanes”. One sample corresponds to one lane.
51. Adjust the width and position of each lane.
52. In the “Bands” tab, add a band for each Lane, and adjust the size of the band in the “Lane Profile” icon so that it includes all the detected GPL stained in blue by the anthrone within a single band.
53. Recover the area under the curve for each band in the “Report” icon and copy them into Excel or any alternative software for quantification and data visualization.

*Note: Results from two independent distinct replicates performed using the manual or semi-automated workflow and subsequently quantified using the ChemiDoc/ImageLab approach are displayed in* Fig 4B*. If some minor changes can be observed in terms of raw values obtained through AUC analysis, they nevertheless demonstrate a great reproducibility with pJL37-mediated GPL inhibition comprised around 76% and 85%*.

#### F. Quantitative imaging and alternative visualization using the open-source software Fiji

54. Open Fiji and load the “Colorimetric Scan” TIFF files acquired using the ChemiDoc Imaging System (Fig 5A).
55. Make sure that the picture of interest is under an 8-bit format. If not, in the “Image” tab, select “Type” and convert the image of interest in 8-bit. This will scale the image to have a range of pixel values comprised between 0 and 255.
56. In the “Edit” tab, invert pixel values using the command “Invert”. This allows the conversion of GPL bands as positive pixels appearing in white on a black background as opposed to black on a white background (Fig 5A).
57. Duplicate this window and use the Look Up Table tab to create a heat-map of pixel intensities that simplify visualization with calibrated colors. In this example, we use the Look Up Table “Fire” (Fig 5A).
58. Use the wand tracing tool to create a region of interest that include the GPL bands of interest. In this example, we drew a rectangle to select the area to be analyzed.
59. Use the shortcut command “Ctrl + t” to include this region of interest in the ROI manager.
60. Use this rectangle and apply it to each sample lane that needs to be analyzed. Further re-adjust the position of region of interest if necessary.
61. Once all the regions of interest have been drawn, make sure to save them using the ROI manager, should you wish to repeat more analysis later on using the same segmented areas.
62. In the ROI Manager, select all the regions of interest, and use the tool “Measure”. This will enable to obtain different type of parameters such as the area analyzed, the min, max, mean, median pixel intensities but also the raw integrated density (RID). To include more parameters, click on the tab “Analyse” and go to the “Set Measurement” Tool.
63. A table of results will appear and can be saved in order to be further processed as mentioned before to enable quantification in R Studio.
64. At this stage, some additional analyses can be performed. Using the ROI Manager and the regions of interest apply the “MultiCrop” function to obtain each area analyzed as an individual picture (Fig 5B).
65. Then, for each of them, use the “Freehand Line” to draw a line that cover the entire area to be analyzed in length.
66. In the “Analyse” Tab, select the “Surface plot” visualization tool. This will create a 2D-surface plot that maps the intensity of each pixel along the axis that has just been created. Save the raw data and generate the plots in R Studio using the ggplot2 package (Fig 5B).
67. Using these data, additional parameters such as the AUC can be obtained using specific packages such as *pracma* (Practical Numerical Math Functions) that enable to perform trapezoidal numerical integration with the *trapz* function [42].
68. A summary of the quantitative results obtained following this analytic workflow is shown in Fig 5C.

## Tips, Tricks and Troubleshooting

In this research protocol, we provide an overview of some simple and efficient assays to evaluate CRISPRi-mediated targeted repression of both essential and non-essential genes in mycobacteria. If CRISPRi has already revolutionized our way of approaching functional genomics in *M. tuberculosis* [43] and other mycobacterial model species [16, 44], our community needs now to further build on this approach to better understand NTM physiology, pathogenesis and drug resistance mechanisms.

**Figure 5.**
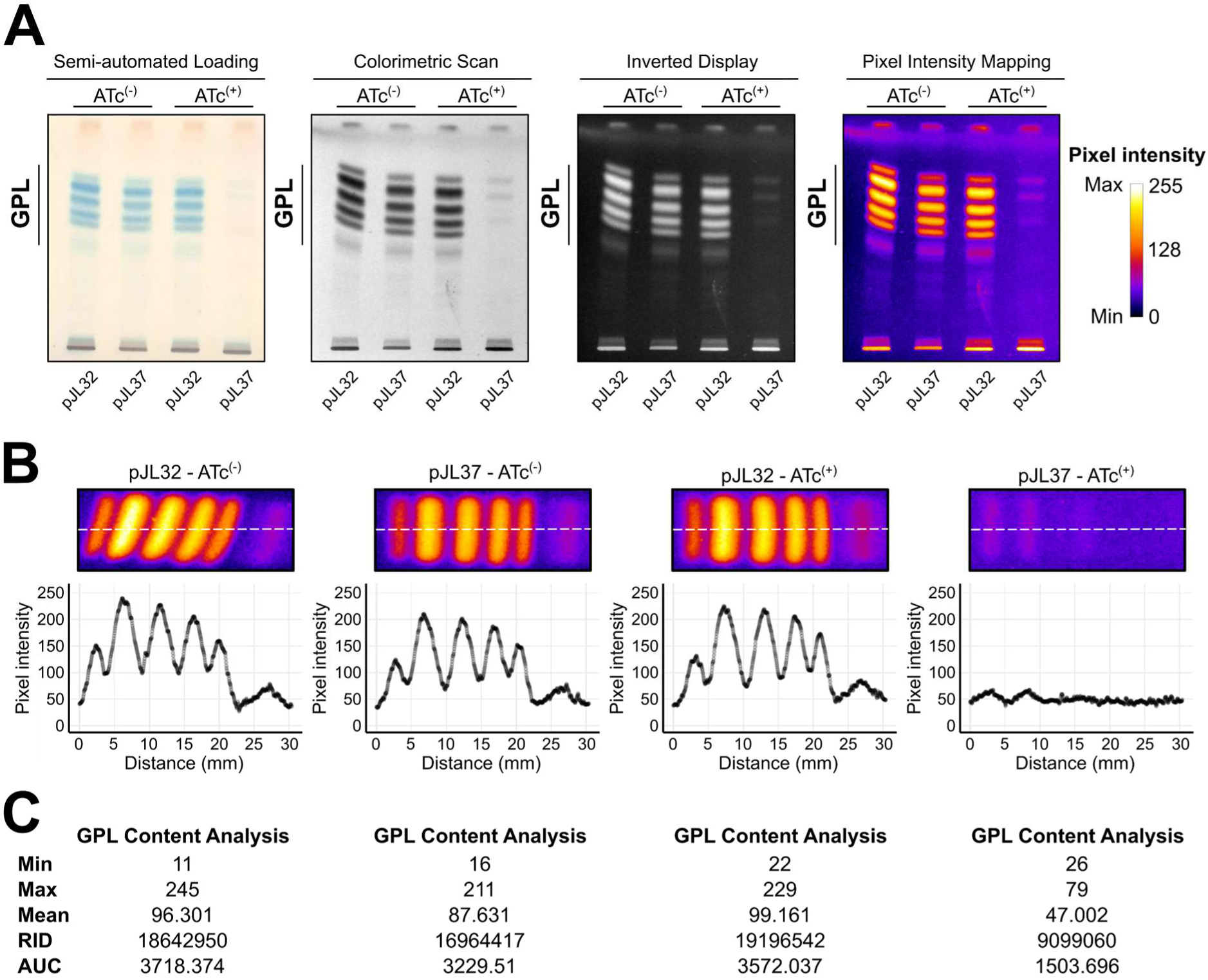
Visualizing and quantifying GPL content using the open-source software Fiji. **(A)** Representative TLC of GPL analysis of *M. smegmatis* mc^2^ 155 pJL32 or pJL37 in the presence or in the absence of ATc. Cells were grown in 7H9 broth ± 100 ng.mL^−1^ of ATc for 48 h. Lipid extraction from normalized dry pellets was analysed by TLC either by performing semi-automated loading or by using a dedicated CAMAG semi-automated equipment. GPL appear in blue upon development, and TLC plates are further scanned in black and white to perform densitometric analyses. In Fiji, images are processed in order to invert pixel intensities which can be subsequently mapped and expressed as color-coded look up table. Pixel intensities range from 0 to 255 units. **(B)** GPL content visualisation and analysis using plot profiling. Pixel intensities are monitored along the x-axis shown in white and expressed in a 0-255 scale. **(C)** Main features regarding the GPL content analysed using Fiji including min, max, mean intensities. Raw integrated density (RID) and area under the curve (AUC) are also displayed.

The different workflows that we report in this manuscript, rely on fast and very affordable microbiological and biochemical methods to analyse specific phenotypes due to targeted genetic repression. Accordingly, these approaches could be implemented virtually in any research groups worldwide. Moreover, by providing simple options that require a minimal set of equipment, such workflows have the potential to be easily set up in BSL2 or BSL3 environments.

Genome-wide analyses have comprehensively listed genes with essential functions in some NTM model species [17, 18, 45]. CRISPRi is among the very few technologies that enables to validate experimentally such findings and further dissect how the repression of such essential gene impacts physiology or any other process of interest. We report two minimalist technics to address whether a gene is essential or not by monitoring mycobacterial growth using conventional broth microdilution or agar spot assays. To do so, we capitalized from previously developed tools that repress the expression of *rpoB* and *mmpL3* [22], two genes encoding major drug targets that are required for bacterial cell division. However, if our experimental setting is simple and enables to rapidly identify whether a gene is required for growth or not, it doesn’t allow, in its current form, to discriminate whether repression of the gene is bacteriostatic or bactericidal. For that, some more advanced time-kill assays must be set-up, as previously reported with the *M. tuberculosis* fumarase (*rv1098c*) [46].

In this protocol, we also demonstrate that the recommended and widely-used concentration of 100 ng.mL^-1^ of ATc exhibits very strong effects in our experimental system and that reducing the ATc doses does not always correlate with an observable decreasing phenotypic response. Indeed, *rpoB* silencing provides the best example for this as all the serial concentrations tested from 100 to 0.04 ng.mL^-1^, all led to growth inhibition in broth medium. Of note, repression of *rpoB* might create a positive loop that enhances the effect of the repression by not only blocking transcription but also depleting the pool of RNA polymerase.

A similar pattern regarding ATc-dependent effects was observed when repressing *mmpL4b* and analysing the remaining GPL content. Indeed, in this experimental setting no major modifications were observed when comparing GPL levels obtained from cells treated with ATc concentrations comprised between 100 and 1 ng.mL^-1^. In this frame, the low levels of detectable GPL suggested that the repression efficacy reached its maximal effect, hence resulting in a plateau. This observation is perfectly in line with previous reports demonstrating that the fine-tuning of knock down efficiency could be performed by targeting different protospacer adjacent motifs and/or modulating the degree of complementarity between the sgRNA and the DNA sequence that is targeted [15, 43]. Then, such strategy could be employed to generate hypomorphs with expression levels ranging from an approximately 200-fold range at a single inducer concentration [15, 47–49]. However, more investigations are required in the near future to fully validate this experimental strategy with an adequate and systematic mapping of transcripts levels.

In our proof of concept, we aimed at targeting GPL translocation as it provides a simple and easy way to assess CRISPRi effectiveness through the S to R morphotype transitioning [30–32]. The mixing procedure presented here, based on the use of two distinct fluorophores, enables to easily detect CRISPRi-mediated effect on mycobacterial colony morphology within the same Petri dish in the absence or the presence of ATc. By doing so, we provide evidences that this is feasible for one gene of interest and could therefore be employed at greater scale to screen genome-wide CRISPRi libraries for some cell-wall alterations and identifying new candidates that are required for cell-wall biosynthesis/maintenance in NTM. Indeed, mapping the genes that are involved in S to R transition, constitute cell-wall associated antigens, virulence factors or play a role in cell-wall integrity is timely and needed as they might represent a new Achille’s heel in mycobacterial species [9].

A known limitation for users while performing CRISPRi are the potential off-targets effects. Several groups have suggested to use additional computational tools to try to predict as much as possible off-target sites while design PAM & sgRNA. Indeed, Rock and colleagues have demonstrated that off-target silencing with the mycobacterial-optimized dCas9Sth1 system is very unlikely [15], therefore performing a thorough *in silico* analysis when designing PAM and sgRNA should prevent any off-target effects. In addition, global transcriptomic or proteomic approaches could be employed to map the biological consequences of the targeted repression and analyse whether they could be due to off-target effects. Overall, the more reliable way of assessing this potential effect remains by performing a complementation strategy where a PAM-mutated version of the gene to be targeted is expressed upon CRISPRi-silencing of the WT copy. By doing so, this additional copy should not be targeted by the sgRNA-dCas9 complex and should therefore fully revert the hypomorph phenotype.

## Abbreviations

NTM: Non-tuberculous mycobacteria
CRISPRi: CRISPR interference
ATc: Anhydrotetracycline
OD_600nm_: Optical density 600nm
GPL: Glycopeptidolipids
TLC: Thin Layer Chromatography.

## Acknowledgements

We would like to acknowledge all members of the Lipolysis and Bacterial Pathogenicity group and the LISM unit for continuous support and insightful discussions. The work developed in our group is supported by the Centre National de la Recherche Scientifique (CNRS) and Aix-Marseille Université (AMU). PS received financial support from the CNRS Biologie, the Agence Nationale de Recherches sur le Sida et les Hépatites virales (ANRS) (project n°ANRS0358), the Agence Nationale de la Recherche (ANR) (ANR-24-CE15-2633) and the French government under the France 2030 investment plan, as part of the Initiative d’Excellence d’Aix-Marseille Université - A*MIDEX and is part of the Institute of Microbiology, Bioenergies and Biotechnology - IM2B (AMX-19-IET-006). JL PhD fellowship was funded by the Ministère de l’Enseignement Supérieur et de la Recherche. WA postdoctoral fellowship was funded by the foundation IHU Méditerranée Infection. PS has also received a FEBS Excellence Award 2023 to support this work and would like to personally thank the FEBS Excellence Awards and Fellowships Office, FEBS Letters and FEBS Open Bio Editorial Offices for their continuous support.

The funders did not play a role in the study design, data collection and analysis, decision to publish, or preparation of the manuscript.

## Authors contribution

PS proposed, conceived and led the study. PS secured funding. SC co-advised the PhD work of JL with PS. SC and CC co-advised the postdoctoral work of WA with PS. VP, WA, JL, ESR, MM and PS performed the experimental work. WA, JL, MM generated the strains and performed CRISPRi-mediated inhibition of essential genes. ESR performed CRISPRi-mediated smooth-to-rough transition experiments. VP performed quantitative analysis of GPL content by TLC. VP, WA, JL, ESR and PS edited figures. All authors provided intellectual input by organizing, analysing and/or discussing data. PS wrote the initial draft of the manuscript with input from VP, JL, and WA. All authors read the manuscript and provided critical feedback before its submission.

## Data accessibility

Original plasmid vectors from the pJL series are available at (https://www.addgene.org/). Any additional data that support the findings of this study are available upon reasonable request from the corresponding author at.

